# TMX5/TXNDC15, a natural trapping mutant of the PDI family is a client of the proteostatic factor ERp44

**DOI:** 10.1101/2024.06.12.598629

**Authors:** Tatiana Soldà, Carmela Galli, Concetta Guerra, Carolin Hoefner, Maurizio Molinari

**Affiliations:** Università della Svizzera italiana, Institute for Research in Biomedicine; CH-6500 Bellinzona, Switzerland; Department of Biology, Swiss Federal Institute of Technology; CH-8093 Zurich, Switzerland; School of Life Sciences, École Polytechnique Fédérale de Lausanne; CH-1015 Lausanne, Switzerland

**Author notes:** Equal contribution (Tatiana Soldà, Carmela Galli).

**Keywords:** Endoplasmic reticulum, ERp44, ERp57, Oxidoreductase, PDI, Protein folding and quality control, TMX5

## Abstract

The endoplasmic reticulum (ER) is the organelle of nucleated cells that produces lipids, sugars and proteins. More than 20 ER-resident members of the Protein Disulfide Isomerase (PDI) family regulate formation, isomerization and disassembly of covalent bonds in newly synthesized polypeptides. The PDI family includes few membrane-bound members. Among these, TMX1, TMX2, TMX3, TMX4 and TMX5 belong to the thioredoxin-related transmembrane (TMX) protein family. TMX5 is the least known member of the family. Here, we establish that TMX5 covalently engages via its active site cysteine residue at position 220 a subset of secretory proteins, mainly single- and multi-pass Golgi-resident polypeptides. TMX5 also interacts non-covalently, and covalently, via non-catalytic cysteine residues, with the PDI family members PDI, ERp57 and ERp44. The association of TMX5 and ERp44 requires formation of a mixed disulfide between the catalytic cysteine residue 29 of ERp44 and the non-catalytic cysteine residues 114 and/or 124 of TMX5 and controls the ER retention of TMX5. Thus, TMX5 belongs to the family of proteins including Ero1α, Ero1β, Prx4, ERAP1, SUMF1 that do not display ER retention sequences and rely on ERp44 engagement for proper inter-compartmental distribution.

## Introduction

The protein disulfide isomerase superfamily includes more than 20 ER-resident members that ensure oxidative folding of newly synthesized polypeptides and participate in clearance of misfolded proteins from the ER lumen (Okumura *et al*, 2015; Patel *et al*, 2020; Pisoni & Molinari, 2016). Few of them, TMX1, TMX2, TMX3, TMX4 and TMX5 are anchored at the ER membrane (Guerra & Molinari, 2020). TMX1 (TXNDC1) is a reductase associated with mitochondria-ER contact sites. It regulates calcium flux and is involved in biogenesis and clearance of membrane-associated polypeptides (Guerra *et al*, 2018; Herrera-Cruz *et al*, 2021; Lynes *et al*, 2012; Matsuo *et al*, 2001; Matsuo *et al*, 2009; Pisoni *et al*, 2015; Raturi *et al*, 2016). TMX2 (TXNDC14) has the active site unconventionally located at the cytoplasmic site of the ER membrane. TMX2 regulates sulfenylation of mitochondrial proteins at mitochondria-ER contact sites to control calcium flux. It also localizes to the nuclear envelope, where it supports nuclear import of select cargo proteins (Chen *et al*, 2024; Guerra & Molinari, 2020; Lynes *et al*., 2012; Oguro & Imaoka, 2019; Quimby & Corbett, 2001). TMX3 (TXNDC10) has a canonical CGHC active site sequence, which corresponds to the catalytic site of PDI. It has oxidase activity in vitro and its b’ domain engages the oxidoreductase’ clients. However, its biological role has not been established (Guerra & Molinari, 2020; Haugstetter *et al*, 2005; Haugstetter *et al*, 2007). TMX4 (TXNDC13) is a reductase located in the ER and in the nuclear envelope, where it regulates the disassembly of disulfide-bonded LINC complexes to promote selective clearance of excess outer nuclear membrane portions during recovery from ER stress (Guerra & Molinari, 2020; Hatahet & Ruddock, 2009; Kucinska *et al*, 2023; Kucinska & Molinari, 2024; Sugiura *et al*, 2010).

TMX5 (TXNDC15) is the least known member of the TMX family. It is a single-span type I protein of 360 amino acids with a large, penta-glycosylated luminal domain and a short C-terminal cytoplasmic tail lacking canonical ER retention signals. TMX5 has 5 cysteine residues (C in single letter code, green, **Fig. 1**). The cysteine at position 220 (red, **Fig. 1**) is embedded in a non-canonical CRFS active site that defines TMX5 as a natural trapping mutant protein (Yang *et al*, 2016). So far, no biological role has been ascribed to TMX5 but mutations in the *TMX5* gene have been associated with the development of the Meckel-Gruber syndrome (MKS), a rare perinatally lethal autosomal recessive disease caused by defective ciliogenesis (Deng & Xie, 2024; Hartill *et al*, 2017; Xu *et al*, 2024).

**Figure 1.**
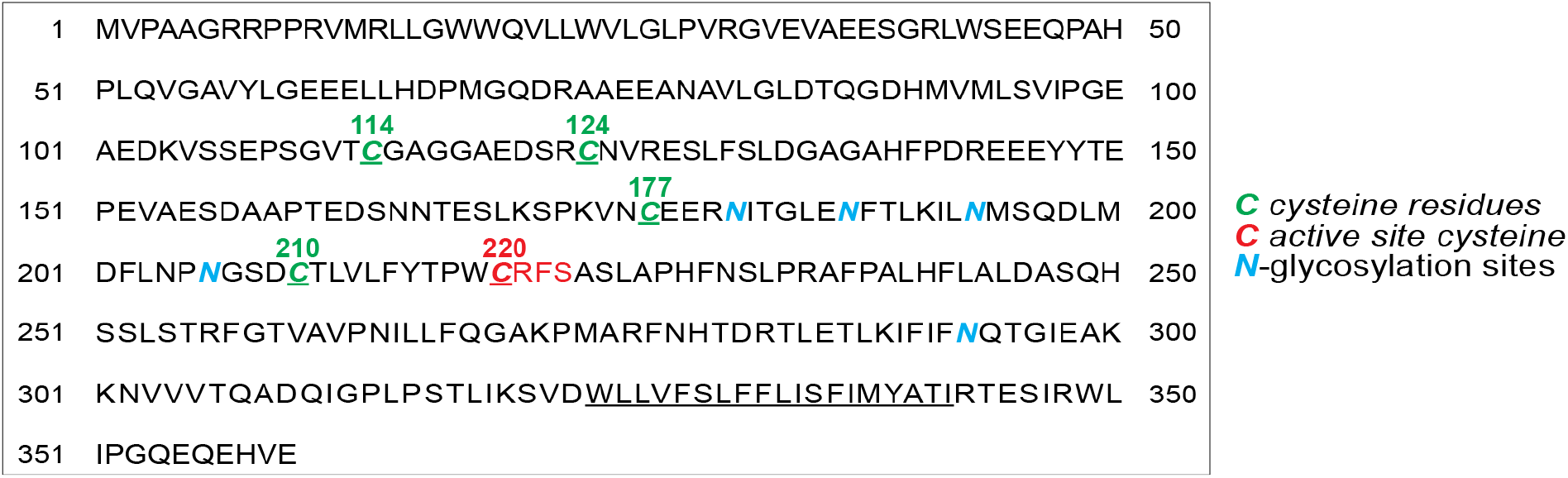
TMX5/TXNDC15 features. TMX5 is a type I glycoprotein (the transmembrane domain is underlined). The CRFS unconventional active site and the active site cysteine residue at position 220 are shown in red. Asparagine residues (N in single letter code, in N-X-S/T consensus sequences for N-glycosylation) are shown in blue. The other 4 cysteine residues at position C114, C124, C177 and C210, whose role is established in this study are shown in green.

Here, we identify the endogenous client proteins trapped in mixed disulfides with the active site cysteine residue 220 of TMX5. We report that endogenous ERp57, PDI and ERp44 associate covalently with non-catalytic TMX5 cysteine residues. We characterize the association between TMX5 and the proteostatic factor ERp44 that controls the ER-to-Golgi cycling of proteins lacking ER retention motifs (Anelli *et al*, 2002). The association involves the TMX5’s cysteine residues 114 and 124 and the ERp44 active site cysteine residue 29 and contributes to the ER retention of TMX5.

## Results and Discussion

### TMX5 localizes in the ER and the Golgi compartments

The presence of 5 N-linked glycans offers the possibility to establish if TMX5 is retained in the ER or traffics to the Golgi compartment by an EndoglycanaseH (EndoH)-sensitivity assay. In fact, upon arrival in the Golgi compartment, resident glycosyl transferases modify the glycoprotein’s N-glycans resulting in an increase of the polypeptide’s apparent molecular weight. Moreover, the N-glycans of proteins retained in the ER are cleaved off by EndoH, whereas the complex glycans generated in the Golgi compartment become resistant to EndoH cleavage (Rothman *et al*, 1984). TMX5 shows a heterogeneous electrophoretic mobility (green line, **Fig. 2A**). This is a symptom of complex glycosylation of TMX5 and indicates that a fraction of the polypeptide is transported to the Golgi compartment. To confirm this, we exposed the immunoisolated TMX5 to EndoH (Rothman *et al*., 1984). These analyses reveal that about 20% of TMX5 displays EndoH-resistant oligosaccharides, thus it reaches the Golgi complex (**Fig. 2B**, lanes 1, 2). In addition to the active site Cys_220_, TMX5 has 4 other cysteine residues at position 114, 124, 177, and 210 (**Figs. 1, 2C**). Individual mutations of Cys_220_, Cys_114_, or Cys_124_ do not affect the cycling of TMX5 between ER and Golgi as shown by the unchanged fraction of complex N-glycans displayed by the proteins (**Fig. 2B**, lanes 3-8). However, mutation of Cys_177_ (lanes 9, 10), or of Cys_210_ (lanes 11, 12), whose proximity in the TMX5 structure hints at their involvement in an intramolecular disulfide bond important to stabilize the TMX5 native structure (**Fig. 2C**), results in efficient retention of the protein in the ER (**Fig. 2B**, lanes 10 and 12). Consistent with the results of the EndoH assay (**Fig. 2B**), the analyses by confocal laser scanning microscopy (CLSM) of the subcellular localization of TMX5 and of the 5 cysteine mutants (**Figs. 2D-2I**) show that TMX5 (**Fig. 2D**), TMX5_C220A_ (**Fig. 2E**), TMX5_C114A_ (**Fig. 2F**), TMX5_C124A_ (**Fig. 2G**) extensively co-localize with the ER marker CNX and with the Golgi marker Giantin. TMX5_C177A_ (**Fig. 2H**) and TMX5_C210A_ (**Fig. 2I**) are retained in the ER, possibly as a consequence of misfolding upon elimination of the structural intramolecular disulfide linking these two cysteine residues.

**Figure 2.**
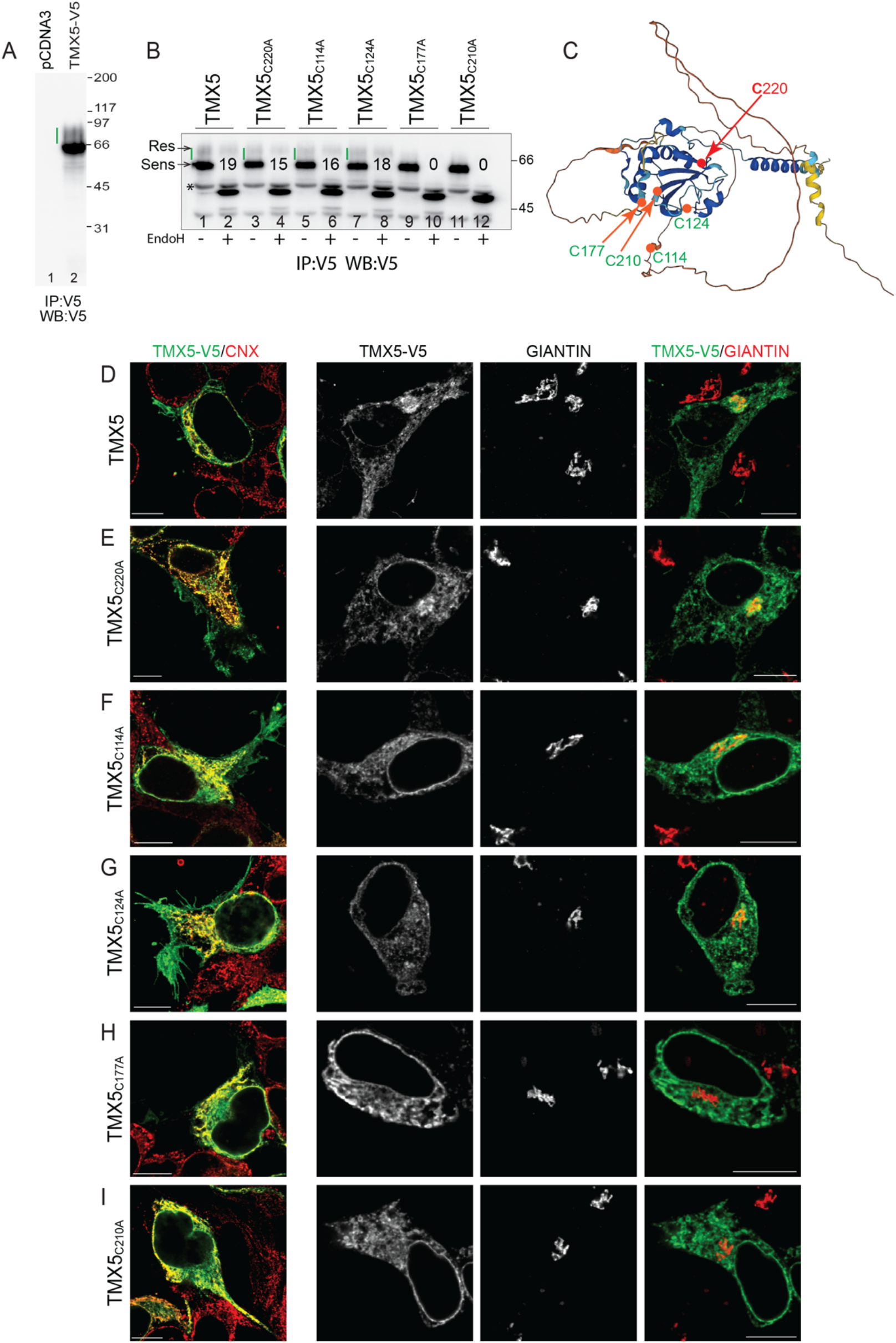
TMX5 localizes in the ER and Golgi. **A** TMX5 separated in a reducing gel (lane 2). The green line shows the complex glycosylated fraction of TMX5 (see panel **B**). **B** Assessment of EndoH-resistant N-glycans of TMX5 and TMX5 cysteine mutants. The % of EndoH-resistant TMX5’s form is given for this experiment. **C** Predicted structure of TMX5, cysteine residues in green, active site cysteine 220 in red. The proximity of cysteine residues 177 and 210 in the TMX5 structure hint at their engagement in an intra-molecular disulfide bond (https://alphafold.ebi.ac.uk/entry/Q96J42). **D** Co-localization in IF of TMX5 with the ER marker CNX and with the trans-Golgi marker Giantin. **E** Same as **D** for TMX5_C220A_. **F** Same as **D** for TMX5_C114A_. **G** Same as **D** for TMX5_C124A_. **H** Same as **D** for TMX5_C177A_. **I** Same as **D** for TMX5_C210A_.

### TMX5 is a natural trapping protein that engages endogenous clients via cysteine 220

We previously assessed the client specificity of the TMX family members *in cellula* (Kucinska *et al*., 2023; Pisoni *et al*., 2015). To this end, we generated the clients-trapping mutant forms of TMX1 (Pisoni et al., 2015), TMX3 and TMX4 (Kucinska et al., 2023) by mutating the C-terminal cysteine of the CXXC active site motif to alanine to substantially delay release of the endogenous clients from the oxidoreductases (Fujimoto *et al*, 2019). This operation was unnecessary for TMX5 that has an unconventional CRFS active site that defines TMX5 as a natural trapping protein (Yang et al., 2016). TMX clients were identified upon co-immunoprecipitation and mass spectrometry (Kucinska *et al*., 2023; Pisoni *et al*., 2015). For TMX5, a V5-tagged version of the protein was expressed in HEK293 cells. After immunoisolation of TMX5 from cell lysates with anti-V5 antibodies, the immunocomplexes were separated in non-reducing (**Fig. 3A**, NRed) and reducing SDS polyacrylamide gels (**Fig. 3A**, Red), which were subsequently silver stained. Disulfide-bonded complexes between endogenous polypeptides and TMX5 are shown in the gel (**Fig. 3A**, NRed, red rectangle) and their identity established by mass spectrometry is listed in **Fig. 3B** (red column). Upon reduction, the disulfide-bonded complexes are disassembled and disappear from this region of the gel (**Figs. 3A**, Red, blue rectangles). Consistently, the TMX5 clients are identified by their disappearance from MS detection when this region of the reducing gel is probed (**Fig. 3B**, blue column). Among the cellular proteins that form mixed disulfides with TMX5, it is worth mentioning the few members of the PDI superfamily (PDI (P4HB), ERp44, ERp57 (PDIA3)), few secretory proteins, and a large spectrum of single and multipass proteins that reside in the membrane of secretory organelles, with preponderance of Golgi-resident polypeptides (**Fig. 3B**). Notably, the endogenous proteins engaging TMX5 are clearly distinct for the endogenous clients of TMX1 (Pisoni et al., 2015), TMX3 and TMX4 (Kucinska et al., 2023). This confirms the exquisite selectivity of individual members of the PDI superfamily, which results from their distinct substrate interaction domains, different co-factors, and different sub-compartmental distribution (Kanemura *et al*, 2020).

**Figure 3.**
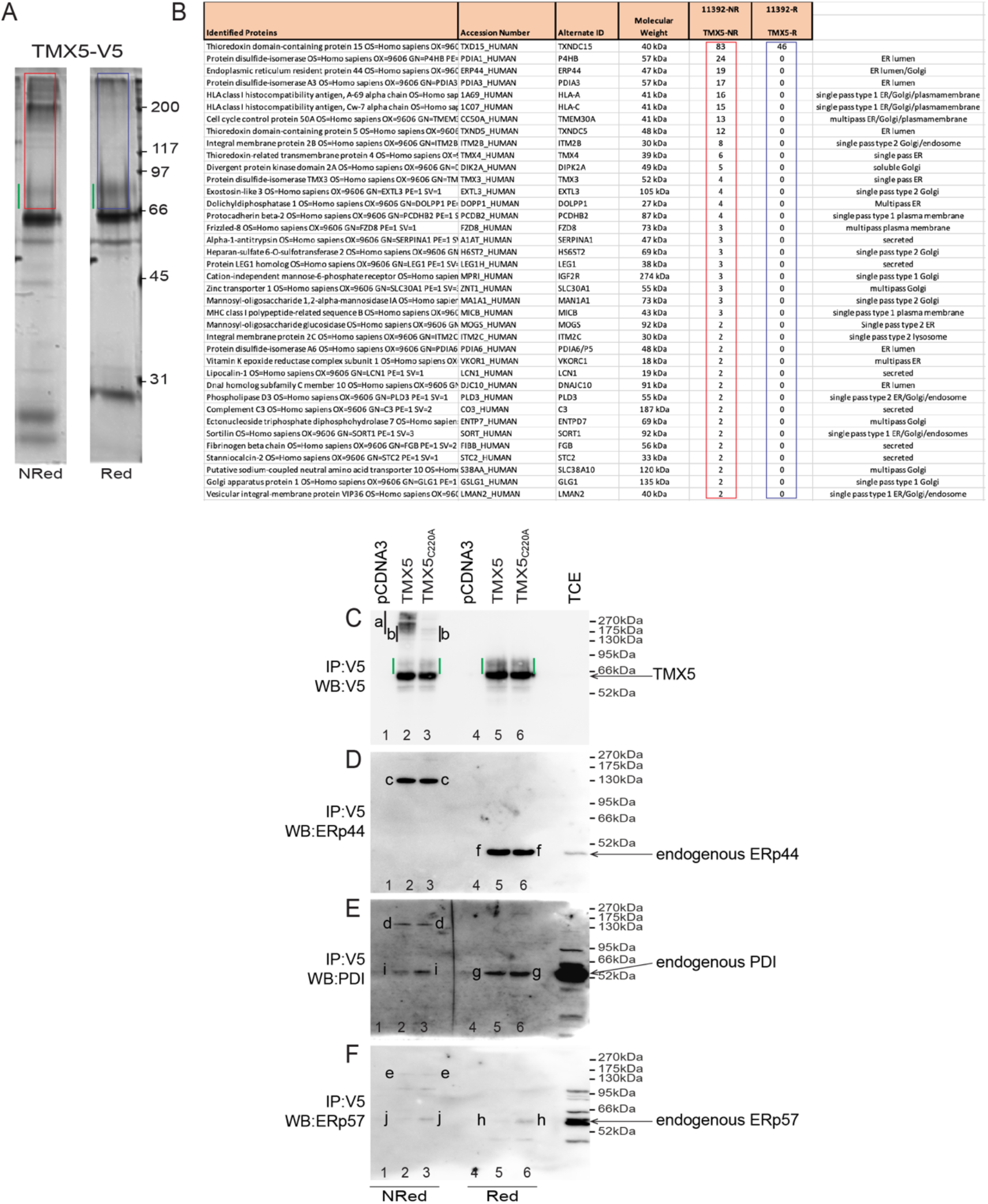
Endogenous interactors of TMX5. **A** V5-tagged TMX5 has been immunoisolated from HEK293 cell lysates. Part of the immunocomplexes were separated in non-reducing/reducing gel and stained with Silver. The endogenous proteins engaged in mixed disulfides with TMX5 (red rectangle) have been identified by LC/MS analyses acer trypsin digestion in gel (two independent experiments). **B** Endogenous polypeptides engaged by TMX5 in mixed disulfides are shown in the red rectangle. **C** The engagement of TMX5 clients in mixed disulfides (lane 2) is substantially inhibited upon replacement of the active site Cys_220_ with alanine (lane 3). *a* shows TMX clients engaged by the active site Cys_220_; *b* shows interaction partners, whose covalent association persists upon mutation of the active site cysteine; green lines show complex glycosylated TMX5. **D** *c* shows the endogenous ERp44 component of TMX5 complexes immunoisolated with anti-V5 antibodies. Upon reduction of the sample, the disulfide bonded complexes are disassembled and the ERp44 component is shown with *f*. ERp44 is engaged in mixed disulfides both with TMX5 (lane 2) and with TMX5_C220A_ (lane 3). **E** Same as **D** for PDI. Note that PDI also associates non-covalently with TMX5 and TMX5_C220A_ (shown with i). **F** Same as **D** for ERp57. Note that ERp57 also associates non-covalently with TMX5 and TMX5_C220A_ (shown with j).

As expected, the replacement of the active site cysteine 220 with an alanine residue substantially hampers the involvement of TMX5 in mixed disulfides (**Fig. 3C**, compare the region shown with *a* in lane 2 *vs*. lane 3). However, we noticed that some covalent interactions were not prevented by this mutation (**Fig. 3C**, shown with *b* in lanes 2 and 3). This shows that select endogenous proteins engage TMX5 by forming mixed disulfides with non-catalytic cysteine residues.

### ERp44 form mixed disulfides engaging the non-catalytic cysteines of TMX5

Analyses of the polypeptides that maintained the covalent association with TMX5_C220A_ (**Fig. 3C**, lane 3, shown with *b*) reveal several members of the PDI superfamily. In fact, both the immunoisolation of TMX5 and of the catalytically inactive TMX5_C220A_ results in the capture of disulfide bonded endogenous ERp44 (**Fig. 3D**, lanes 2-3, *c*), of disulfide-bonded endogenous PDI (**Fig. 3E**, lanes 2-3, *d*), of disulfide bonded endogenous ERp57 (**Fig. 3F**, lanes 2-3, *e*). These covalent complexes disappear upon reduction (**Figs. 3D-3F**, lanes 5-6) and the signals collapse into the monomeric forms of ERp44 (**Fig. 3D**, *f*), PDI (**Fig. 3E**, *g*) or ERp57 (**Fig. 3F**, *h*). PDI and ERp57 also associate non-covalently with TMX5 as shown by their monomeric forms that co-precipitate both with TMX5 and with the catalytically inactive TMX5_C220A_ (shown with *i* and *j*, respectively in **Figs. 3E, 3F**, lanes 2-3). These data reveal that PDI, ERp57 and ERp44 are not clients of TMX5. Rather, they could play a role in ruling the biological activity of TMX5 by acting as co-factors. PDI and ERp57 oxidoreductases could regulate the release of clients from TMX5. In fact, the N-terminal cysteine of the TMX5 active site can nucleophilically attack a free thiol group in protein substrates, but the interactions can be resolved only by the intervention of an external cysteine provided by glutathione, or by another PDI (Yang *et al*., 2016). More interesting in the context of our study is the covalent association of TMX5 with ERp44, an unconventional member of the PDI superfamily previously reported to regulate the positioning in the secretory line of proteins lacking the ER retrieval sequence (Otsu *et al*, 2006; Tempio & Anelli, 2020), a feature that TMX5 shares with a number of polypeptides acting in early compartments of the secretory line.

### The covalent association of ERp44 with TMX5 relies on ERp44’s cysteine residue 29

Notably, complex glycosylation is not observed for TMX3 and TMX4 that, in contrast to TMX5, possess C-terminal KKXX retention sequences that efficiently retains them in the ER or in the contiguous nuclear envelope membrane (Guerra *et al*., 2018; Guerra & Molinari, 2020; Kucinska *et al*., 2023; Pisoni *et al*., 2015). In this context, the finding that endogenous ERp44 is a major interacting partner of TMX5 (**Figs. 3B, 3D**) is relevant. In fact, ERp44 cycles between ER and Golgi to ensure the retention of key factors designed to act in the ER but devoid of suitable localization motifs including Ero1α, Ero1β, Prx4, ERAP1, SUMF1 (Fraldi *et al*, 2008). The ER retention function of ERp44 is exerted by interacting with clients via the catalytic site cysteine residue at position 29 (Tempio & Anelli, 2020). To verify whether the active site Cys_29_ of ERp44 is involved in the association with TMX5, HA-tagged ERp44 or ERp44_C29S_ were co-expressed in HEK293 cells with V5-tagged TMX5. Complexes were immunoisolated from detergent lysates with anti-V5 antibodies and separated under non reducing or reducing conditions to verify the presence of HA-tagged ERp44 and ERp44_C29S_ in the immunocomplexes by WB (**Figs. 4A, 4B**, respectively). Analyses of the WB confirm that ERp44 engages TMX5 in mixed disulfides (shown with *a*, lane 2, **Fig. 4A**), whereas ERp44_C29S_ does so with much reduced magnitude (lane 3).

**Figure 4.**
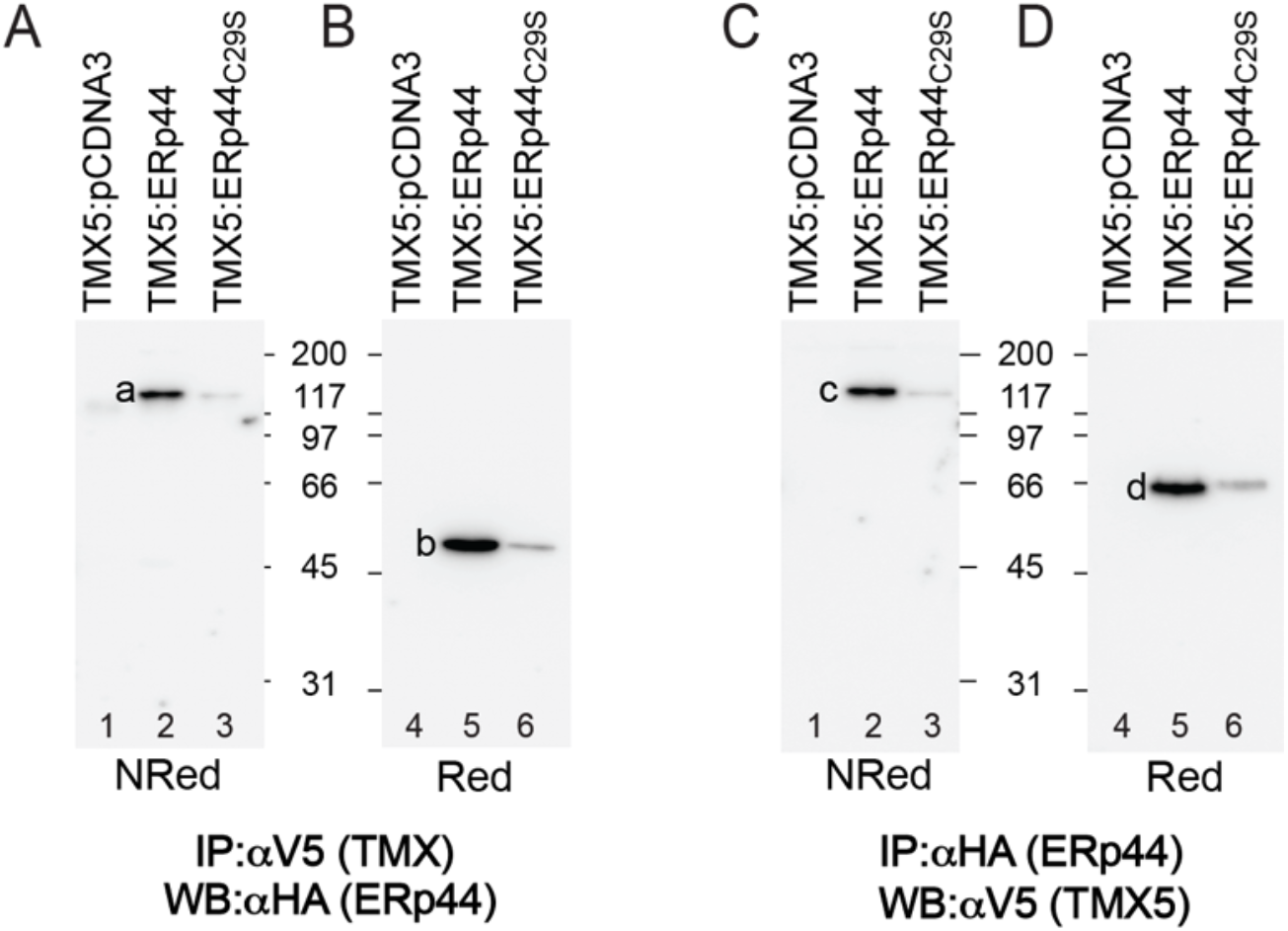
TMX5 engages ERp44 in mixed disulfides via its catalytic Cys_29_. **A** Formation of mixed disulfides between TMX5 and ERp44 (lane 2) or ERp44_C29S_ (lane 3). Mixed disulfides are immunoisolated with anti-V5 antibodies, are separated in nonreducing SDS-PAGE. The ERp44 component (indicated with *a*) is seen by WB with an anti HA antibody. **B** Same as **A** , but the mixed disulfides are disassembled by running the gel under reducing conditions. *b* shows ERp44. **C** Same as **A** where the mixed disulfides have been immunoisolated from detergent extract with an anti-HA antibody and the TMX5 component is revealed in WB with an anti-V5 antibody (indicated with *c*). **D** Same as **B** for the TMX5 (indicated with *d*).

Disassembly of the complexes under reducing conditions reveals the ERp44 component of the mixed disulfides (shown with *b*, lane 5, **Fig 4B**). The composition of the complexes was confirmed by their immunoisolation with an anti-HA antibody to capture the ERp44 component, and detection of the TMX5 component in WB with an anti-V5 antibody (**Figs. 4C, 4D**, lanes 2 and 5). Notably, the TMX5 that co-precipitates with ERp44 (**Fig. 4D**, lane 5, shown with d) lacks the slow migrating, complex glycosylated polypeptide fraction that cycles in the Golgi compartment shown with green lines in **Figs. 2A, 2B, 3A, 3C**. Thus, ERp44 associates covalently via the active site Cys_29_ with the population of TMX5 located in the ER and prevents its delivery to the Golgi compartment.

### The covalent association of ERp44 with TMX5 relies on TMX5’s cysteine residues 114 and 124

Mutation of the catalytic site Cys_220_ of TMX5 does not inhibit the covalent association of endogenous ERp44 (**Fig. 3D**). To establish the TMX5 cysteine residue(s) involved in the mixed disulfide with ERp44, HA-tagged ERp44 was co-expressed with individual V5-tagged cysteine mutants of TMX5 in HEK293 cells.

The formation of covalent complexes between ERp44 and TMX5 variants was tested upon immunoisolation of the complexes with anti-HA antibodies, separation of the immunocomplexes in non-reducing/reducing WB and detection of the TMX5 interacting partner with anti-V5 antibodies. As shown in **Fig. 3D**, ERp44 engages in mixed disulfides with TMX5 (**Fig. 5**, lane 1) and TMX5_C220A_ (**Fig. 5**, lane 2). The mutation of Cys_114_ and of Cys_124_ to alanine residues prevents covalent engagement of the TMX5 protein by the active site cysteine of ERp44 (lanes 3, 4). Mutations of Cys_177_ (lane 5) and of Cys_210_ (lane 6) allows the covalent engagement of TMX5 with formation of aberrant complexes possibly resulting from TMX5 misfolding/instability due to deletion of the structural disulfide that binds these two cysteine residues (red lines, lanes 5, 6). Thus, the active site Cys_29_ of ERp44 engages Cys_114_ and Cys_124_ of TMX5 in mixed disulfides.

**Figure 5.**
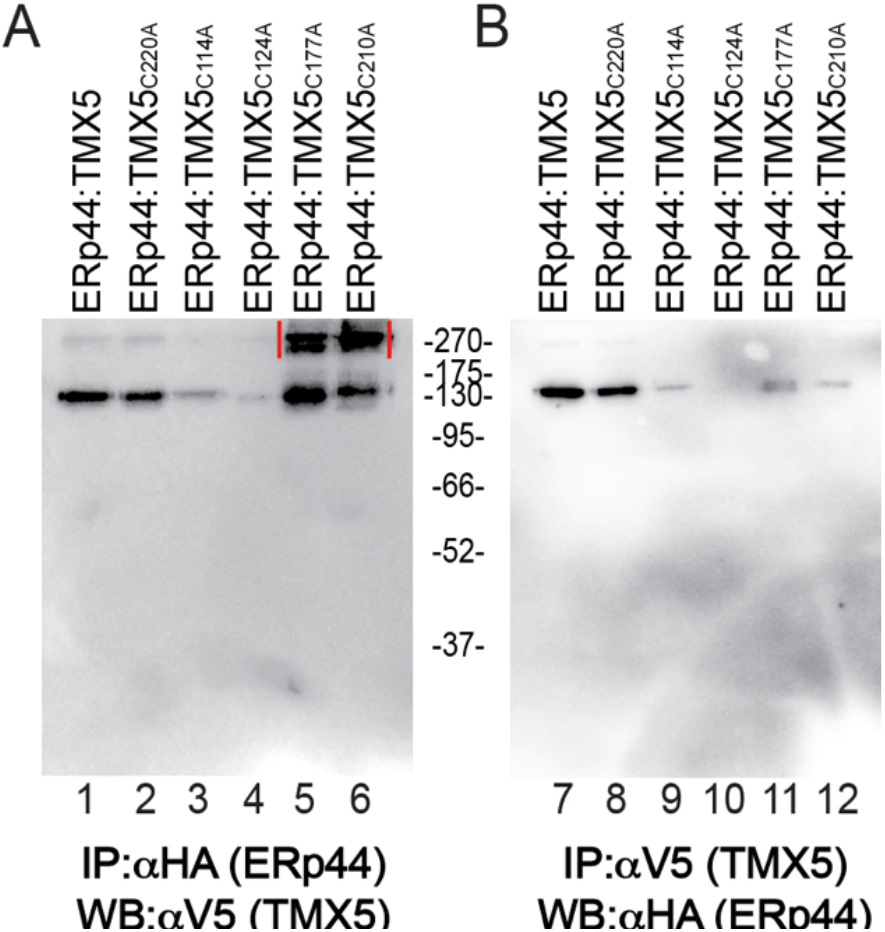
ERp44 engages Cys_114_ and/or Cys_124_ of TMX5 in mixed disulfides. **A** Co-precipitation of cysteine mutants of TMX5 with ERp44. The mixed disulfides are separated in a non-reducing gel. Here the TMX5 component of the ERp44-S-S-TMX5 complex is shown. **B** Co-precipitation of ERp44 with TMX5. The ERp44 component of the ERp44-S-S-TMX5 complex is shown in the non-reducing gel.

### The association with ERp44 retains TMX5 in the ER and reduces engagement of endogenous clients

The co-expression of active ERp44 substantially reduces the fraction of TMX5 engaged with endogenous clients in mixed disulfides (compare **Fig. 6A**, lanes 1 *vs*. 2, shown with *a*). The co-expression of the ERp44_C29S_ mutant that does not engage TMX5 in covalent complexes (**Fig. 4**), does not affect the formation of mixed disulfides between TMX5 and its clients (**Fig. 6A**, lane 3).

**Figure 6.**
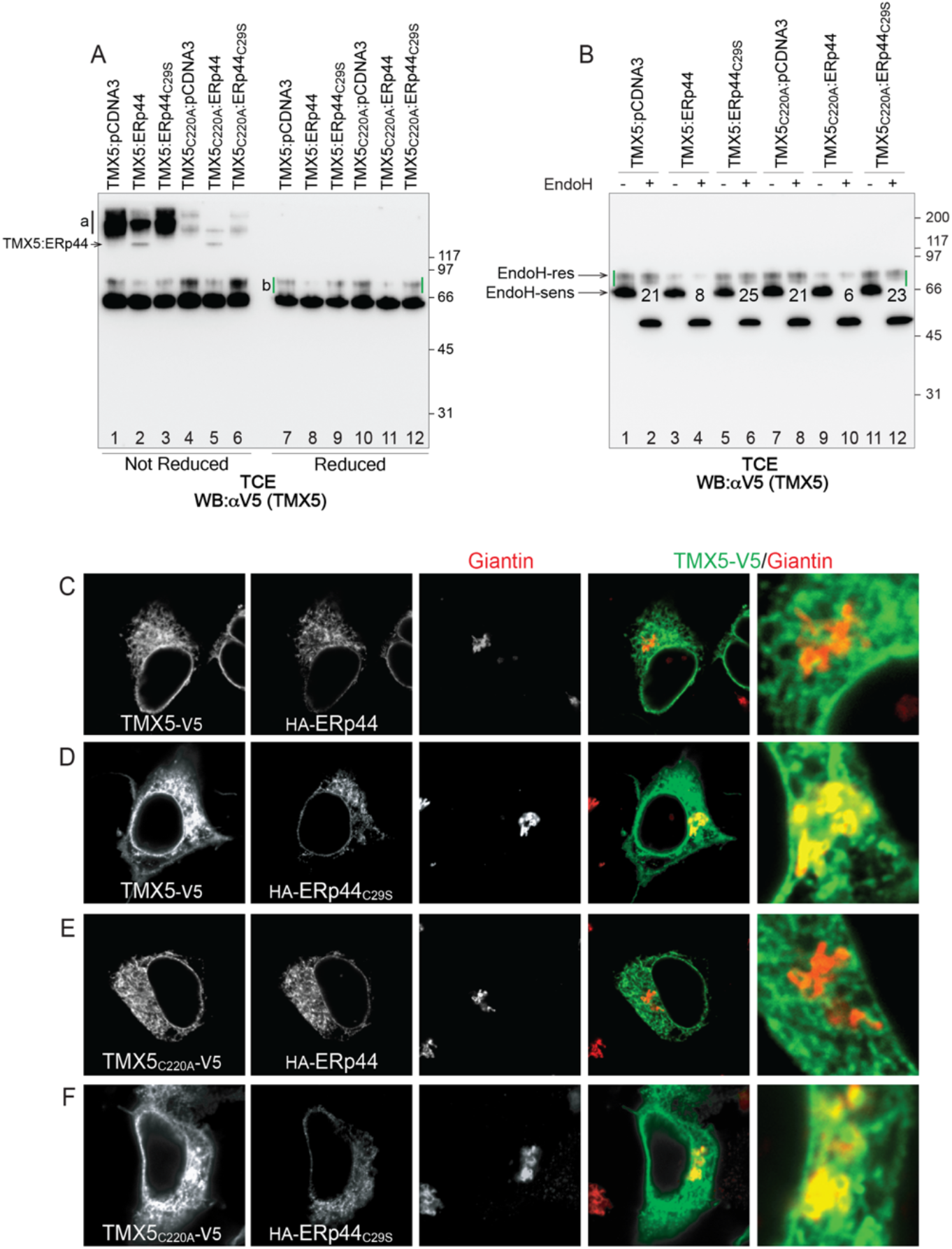
The association with ERp44 retains TMX5 in the ER and reduces engagement of endogenous clients. **A** Western blot analysis of post nuclear fractions of HEK293 cells co-transfected with V5-tagged TMX5 and HA-tagged ERp44 or ERp44_C29S_. The TMX component of the complexes is revealed with anti-V5 antibody in non-reducing and reducing conditions (lanes 1-6 and 7-12, respectively). Mixed disulfides between TMX5 and endogenous clients are shown with the black line *a*. TMX5:ERp44 mixed disulfides are shown in lanes 2 and 5. The complex glycosylated, Golgi form of TMX5 is shown with the green lines *b*. **B** EndoH assay of the same samples. The EndoH-resistant, Golgi forms of TMX5 are shown with green lines. The % of EndoH-resistant TMX5’s form is given for this experiment. **C** Lack of co-localization of TMX5 with the Golgi marker Giantin upon co-expression of active ERp44. **D** Co-localization of a fraction of TMX5 with Giantin upon co-expression of inactive ERp44_C29S_. **E** Same as **C** for inactive TMX5_C220A_. **F** Same as **D** for inactive TMX5_C220A_ showing lack of co-localization with Giantin upon co-expression of ERp44_C29S_.

Analyses of the electrophoretic mobility of TMX5 (**Fig. 6A**, lanes 7-12, green lines) and the EndoH assay (**Fig. 6B**) also reveal that the association with ERp44 retains TMX5 in the ER as shown by the substantial reduction in TMX5 complex glycosylation from about 20% EndoH-resistant fraction in mock-transfected cells (**Figs. 2B, 6B**, lanes 1, 2) and in cells expressing ERp44_C29S_ (**Fig. 6B**, lanes 5, 6) to less than 10% (**Fig. 6B**, lanes 3, 4). Replacement of the active site TMX5 cysteine 220 with alanine, abolishes the engagement of clients in mixed disulfides (**Figs. 3C, 6A**, lanes 4-6). This mutation does not affect the fraction of TMX5 that acquires EndoH-resistant N-glycans as symptom of ER to Golgi trafficking (**Figs. 6B**, lane 7, 8), does not prevent engagement of ERp44 (**Fig. 3D**), nor the capacity of ERp44 to inhibit release of TMX5 from the ER (**Figs. 6B**, lanes 9-12). The role of ERp44 in determining the intracellular localization of TMX5 has been confirmed by CLSM showing that TMX5 and TMX5_C220A_ are excluded from the Golgi compartment when co-expressed with active ERp44 (**Figs. 6C, 6E**), but traffic to the Golgi compartment when co-expressed with the active site mutant form ERp44_C29S_ (**Figs. 6D, 6F**).

All in all, our study offers a preliminary set of information on a neglected member of the PDI family by showing that TMX5 engages a subset of cellular proteins in mixed disulfide via the active site cysteine 220. TMX5 clients are clearly distinct from the clients of TMX1 (Pisoni et al., 2015), TMX3 and TMX4 (Kucinska et al., 2023) confirms the selectivity of individual members of the PDI superfamily, which results from their distinct substrate interaction domains, different co-factors, and different sub-compartmental distribution (Kanemura *et al*., 2020). Unfortunately, our attempts to silence the expression of TMX5 failed, and we were therefore unable to evaluate the consequences of lack of TMX5 on the client’s fate. Our results allow to add TMX5 to the limited group of proteins including Ero1α, Ero1β, Prx4, ERAP1, SUMF1 that lack the ER retrieval sequence and rely on ERp44 engagement to control their sub-cellular localization. Notably, in contrast to the other ERp44 clients, TMX5 is tethered at the ER membrane. Why these proteins that exert most of their known activities in the ER do not rely on conventional C-terminal retention/retrieval sequences to be retained in the compartment is a matter of discussion, but it is likely that under specific circumstances, they are called to exerts action in the Golgi compartment, at the cell surface, or extracellularly (Tempio & Anelli, 2020).

## Materials and Methods

### Cell Culture, Transient Transfection

Human Embryonic Kidney 293 (HEK293) and Mouse Embryonic Fibroblasts (MEF) cell lines were cultured at 37°C and 5% CO2 in Dulbecco’s Modified Eagle Medium (DMEM) with high glucose (GlutaMAX™, Gibco) supplemented with 10% Fetal bovine serum (FBS, Gibco). Transient transfections were carried out using JetPrime (Polypus) in DMEM 10% FBS supplemented with non-essential amino acids (NEAA, Gibco) following the manufacturer’s protocol. Experiments were performed 17h after transfection.

### Antibodies and expression plasmids

S1 Table: details of antibodies used. Plasmids encoding for TMX5-V5 WT and cysteine mutants were synthesized by Genscript in pUC57 and subcloned in pcDNA3.1(-) through restriction digestion of BamHI and HindIII (NEB) sites flanking the target sequence. Ligation was carried out using T4 DNA ligase (NEB) at a 3:1 insert to vector ratio, following the manufacturer’s protocol. Plasmids were subsequently amplified and isolated from JM109 bacteria (Promega) using the GenElute™ HP plasmid MidiPrep kit (Sigma). Plasmids for expression of ERp44wt and C29S mutant were a kind gift of Roberto Sitia.

### Cell lysis, Immunoprecipitation (IP), Western Blot (WB) and EndoH treatment

HEK293 cells were washed with ice-cold 1xPBS containing 20 mM N-ethylmaleimide (NEM), then lysed with RIPA buffer containing 20 mM NEM and protease inhibitors (1 mM phenylmethylsulfonyl fluoride (PMSF), 16.5 mM Chymostatin, 23.4 mM Leupeptin, 16.6 mM Antipain, 14.6 mM Pepstatin) for 20 min on ice. Lysates were subjected to centrifugation for 10 min at 4°C, 10,600 g to extract the post-nuclear supernatant (PNS). For immunoprecipitation PNS were incubated with Anti-V5 Agarose Affinity Gel conjugated beads or with Protein-A beads (Sigma, 1:10 w/v swollen in PBS) and anti-HA antibody for 2 h at 4°C. Samples were than washed with 1% Triton in HBS 1x pH 7.4 and then dried. Dried beads were resuspended in sample buffer and boiled at 95°C for 5 min. After boiling, samples without (Non reducing) or with 100mM DTT (Roche)(reducing) were subjected to SDS-PAGE. For detection of proteins by Western Blot (WB) protein bands (IP or TCE) were transferred from polyacrylamide gel to a PVDF membrane using a TransBlot Turbo device (BioRad). PVDF membrane was blocked for 10 min with 8% blocking milk in Tris-buffered saline, 0.1% Tween 20 (v/v) (TBS-T) and incubated with primary antibodies (Table **S1**) with shaking. After primary antibody washout with TBS-T, membranes were incubated with HRP-conjugated secondary antibody or with HRP-conjugated protein A (Table **S1**) for 45 min at RT while shaking. Protein bands were detected on a Fusion FX7 chemiluminescence detection system (Vilber) using WesternBright™ Quantum or WesternBright ECL HRP Substrate (Witec AG) following the manufacturer’s protocol. Protein bands were quantified with FIJI/ImageJ software. For EndoH (NEB) treatment proteins from immunoprecipitated samples or from TCE were split into two aliquots and incubated in the presence or absence of 5mU of EndoH for 3 h at 37 °C according to the manufacturer’s protocol. Samples were then analyzed by SDS-PAGE.

### Mass spectrometry

HEK cells, transfected with an empty vector or transfected with V5-tagged TMX5 were rinsed with PBS and 20 mM NEM. Cells were lysed with RIPA buffer supplemented with 20 mM NEM and protease inhibitors for 20 min on ice. PNS Immunoisolated with anti V5 Agarose beads were washed three times with 1% Triton in HBS 1x pH 7.4 and then dried. Mass spectrometry analysis was performed at the Protein Analysis Facility, University of Lausanne, Lausanne, Switzerland.

### Confocal laser scanning microscopy (CLSM)

HEK cells were seeded on poly-L coated (Sigma) glass coverslips (VWR) and transiently transfected as indicated using JetPrime (Polypus) in DMEM 10% FBS supplemented with NEAA (Gibco). Seventeen hours after transfection, cells were fixed at RT for 20 min in 3.7% formaldehyde (FA, v/v) in PBS. Coverslips were incubated

for 20 min in permeabilization solution (PS, 10 mM HEPES, 15 mM glycine, 10% goat serum (v/v), 0.05% saponin (w/v)). Following permeabilization, primary antibodies (Table S1) diluted 1:100 in PS were applied for 2 h, washed three times in PS, and then incubated with Alexa Fluor-conjugated secondary antibodies diluted 1:300 in PS for 45 min. Cells were washed three times with PS and once with deionized water and mounted on a drop of Vectashield (Vector Laboratories) supplemented with 40,6-diamidino-2-phenylindole (DAPI). Coverslips were imaged using a Leica TCS SP5, microscope equipped with Leica HCX PL APO lambda blue 63.0 × 1.40 oil objective. Leica LAS X software was used for image acquisition, with excitation provided by 488-, 561-, and 633-nm lasers and fluorescence light was collected within the ranges of 504-587 nm (AlexaFluor488), 557-663 nm (TMR) and 658-750 nm (AlexaFluor646), respectively with pinhole 1 AU. Image post-processing was performed with Adobe Photoshop.

---

**S1 Table.** Details of antibodies used.

| Antibody | Manufacturer | Catalog number | Dilution or Concentration |
| --- | --- | --- | --- |
| Mouse anti-V5 | Thermo Fisher Scientific | R96025 | 1:5000 (IB); 1:100 (CLSM) |
| Rabbit anti-HA | Sigma | H6908 | 1:3000 (IB); 1:100 (CLSM) |
| Anti-V5 Agarose Affinity Gel antibody | Sigma | A7345 |  |
| Rabbit anti-CNX | Kind gift from A. Helenius | Not applicable | 1:3000 (IB) |
| Mouse anti-GAPDH | Merck | MAB374 | 1:30000 (IB) |
| Rabbit anti-ERp44 | Kind gift from R. Sitia |  | 1:1000 (IB) |
| Rabbit anti-PDI | Stessgen | SPA890 | 1:1000 (IB) |
| Rabbit anti-ERp57 | Kind gift from G. Hämmerling |  | 1:1000 (IB) |
| Rabbit anti-Giantin | Biolegend | 920342 | 1:100 (CLSM) |
| Protein A HRP-conjugated | Invitrogen | 101023 | 1:20000 (IB) |
| Goat anti-mouse HRP-conjugated | Southern Biotech | 1031-05 | 1:20000 (IB) |

